# Proteomic analysis to classify extracellular vesicles and plasmodesmata in Arabidopsis cell culture supernatants

**DOI:** 10.1101/2024.05.17.594688

**Authors:** Takahito Takei, Teppei Asou, Nobuteru Hamakawa, Sho Tanaka, Ryo Noguchi, Kohei Sumiyoshi, Kenji Sueyoshi, Koshi Imami, Takahiro Hamada

**Affiliations:** Department of Bioscience, Faculty of Life Science, Okayama University of Science, Ridaicho 1-1, Okayama, 700-0005, Japan; Department of Applied Chemistry, Graduate School of Engineering, Osaka Metropolitan University, Osaka, 599-8531, Japan; Department of Chemistry, School of Science, Kitasato University, Sagamihara, Kanagawa, 252–0373, Japan; Proteome Homeostasis Research Unit, RIKEN Center for Integrative Medical Sciences, Tsurumi-ku, Yokohama, Kanagawa 230-0045, Japan

## Abstract

The plant apoplast contains diverse proteins. In this study, we analyzed the extracellular vesicle proteome using cell culture supernatants. Unexpectedly, many plasmodesmata proteins were identified in the extracellular vesicle fraction. The comparison between the cell culture EV proteome and the plasmodesmata proteome revealed that EV and plasmodesmata proteins were indistinguishable. Highly sensitive imaging analyses showed that PDLP3-GFP, a representative plasmodesmata marker, was localized to plasmodesmata, but it was also present as small granules in apoplast regions. Next, we attempted to distinguish EV proteins from plasmodesmata proteins. The step-wise density gradient ultracentrifugation and clustering analysis separated EV protein clusters (represented by tetraspanins and membrane trafficking proteins) from plasmodesmata clusters (represented by PDLPs and PDCBs). These results provide the basis for future explorations of plant plasmodesmata and EVs.

## Introduction

Extracellular vesicles (EVs) are small membrane-enclosed particles found in the apoplast regions. Very little is known about how plant cells secrete EVs, but, it appears that multi-vesicular body (MVB) is a major source. The main function of MVBs is to degradate proteins and lipids by transporting them into the vacuole. For example, proteins ubiquitinated in the plasma membrane are transported to MVB via endocytosis, after which the MVB fuse to the vacuolar membrane (Mosesso et al., 2019). In some cases, MVBs are bound to the plasma membrane and secrete EVs. Several SNAREs and Rab GTPases have been identified for the fusion of MVBs to the plasma and vacuolar membranes (Ito and Uemura, 2022). For example, ARA6, VAMP721/VAMP722 and SYP121/SYP122 mediate the fusion of MVBs to the plasma membrane, whereas ARA7/RHA1 and VAMP727 are involved in the fusion of MVBs to the vacuolar membrane.

However, EVs are not only derived from MVBs, suggestive of a relatively complex process. The secretion of EVs is activated in plants infected by pathogenic fungi and bacteria (Wang et al., 2017; Hansen and Nielsen, 2017). During an infection by pathogenic fungi, vacuolar and plasma membranes fuse together, which leads to the extracellular secretion of vacuolar components, including MVB-derived vesicles and autophagosomes (Cui et al., 2020). Although *Arabidopsis thaliana* (Arabidopsis) plants are normally resistant to *Golovinomyces orontii*, a mutation to *PEN1* (*SYP121*) results in plants that are susceptible to this phytopathogenic fungus that causes powdery mildew (Collins et al., 2003; Assaad et al., 2004; Lipka et al., 2005). The *SYP121* gene encodes a SNARE protein involved in fusing membranes, suggesting it contributes to the secretion of EVs. However, SYP121 was also detected in EVs, implying it may not be involved in the fusion of MVBs to the plasma membrane (Meyer et al., 2009; Nielsen and Thordal-Christensen, 2012). These findings reflect the importance of membrane trafficking factors for plant responses to disease, but the precise functions of SYP121 in plant resistance to phytopathogenic fungi remain unclear.

Identifying the contents of purified EVs is critical for elucidating EV origins and functions. The purification of EVs requires the recovery of apoplasts that are not contaminated with the inner cellular components and vascular components. Obtaining apoplasts from the imbibed leaf lumen via centrifugation is reportedly the optimal method (Rutter et al., 2017). In this method, the apoplast recovery solution is infiltrated into the leaf lumen and then gently recovered by a slow-speed centrifugation. The apoplasts are subsequently collected following a density gradient centrifugation. The use of this method resulted in the identification of 598 proteins in the EV fraction of Arabidopsis (Rutter and Innes, 2017). Among these proteins, several EV markers, including tetraspanins and SYP121, were identified, indicative of the successful recovery of EVs. Additionally, biotic and abiotic stress-responsive proteins, membrane trafficking proteins, plasma membrane-localized proteins, and vacuolar proteins were enriched. Moreover, several ribosomal proteins, cytoplasmic enzymes, and cell wall-associated proteins were also identified. Similarly, 237 proteins were identified in the EV fraction of sunflower seedlings (Regente et al., 2017). The protein composition did not differ significantly from that of the Arabidopsis EV fraction. The increase in EV secretion following an infection by pathogenic fungi and bacteria may be associated with a change in the EV protein composition (e.g., increase in the abundance of resistance-related proteins). However, a previous study on Arabidopsis showed the EV contents are not significantly affected by a *Pseudomonas syringae* infection (Rutter and Innes, 2017). An in depth analysis of the EV proteome of *Botrytis cinerea*-infected Arabidopsis identified 981 proteins including several RNA-binding proteins (e.g., AGO1 and the DEAD-box RNA helicases RH11 and RH37) (He et al., 2021).

In this study, we conducted a proteomic analysis of Arabidopsis EVs obtained from cell culture supernatants. The protein composition of the EV proteome was similar to that of plasmodesmata (PD) proteomes (Fernandez-Calvino et al., 2011; Brault et al., 2019). Confocal microscopy images revealed that a typical PD marker, PDLP3-GFP, was localized to small particles in the apoplast region where PDs are absent. The optimized iodixanol density gradient ultracentrifugation proteomic analysis and the in depth clustering analysis enabled the separation of EV and PD proteins. The apoplast-secreted EV- and PD-related proteins newly identified in this study will be useful for future research on plant apoplasts as well as investigations involving EV and PD.

## Results

### Extracellular vesicles were detected in the plant cell culture medium

The supernatant of a plant cell culture is ideal biochemical materials for plant apoplast studies. More specifically, it is highly homogeneous and can be obtained from large cell cultures relatively easily. Thus, to study plant EVs, we focused on cell culture supernatants. First, to verify the presence of EVs in plant cell culture supernatants, the cell culture supernatant of tobacco BY-2 cells or Arabidopsis MM2d cells were prepared via centrifugation for the subsequent analysis (Fig. 1A). The sedimented fraction on the Percoll cushion was stained with BODIPY TR Ceramide to detect sphingolipids, which are abundant in animal EVs. Many particles were detected only in the sedimented fraction on the Percoll cushion (Fig. 1B, C and D, Supplemental Movie 1). To analyze these sedimented particles, the size and number of particles were measured using qNano. The results showed that the size of these particles ranged from 70 to130 nm with a peak at 87.5 nm (Fig. 1E). The size distribution of these particles is similar to that of typical exosome like EVs (Meldolesi 2018). Furthermore, to check whether EVs were collected in the sedimented fraction, we used green fluorescent protein (GFP)-tagged SYP121, which is a plant EV marker (He et al., 2021; Rutter and Innes, 2017). The SYP121-GFP signal was detected in the sedimented fraction on the Percoll cushion and overlapped with BODIPY TR Ceramide staining (Fig. 1F). These results indicated that EVs obtained from plant cell culture supernatants.

**Figure 1.**
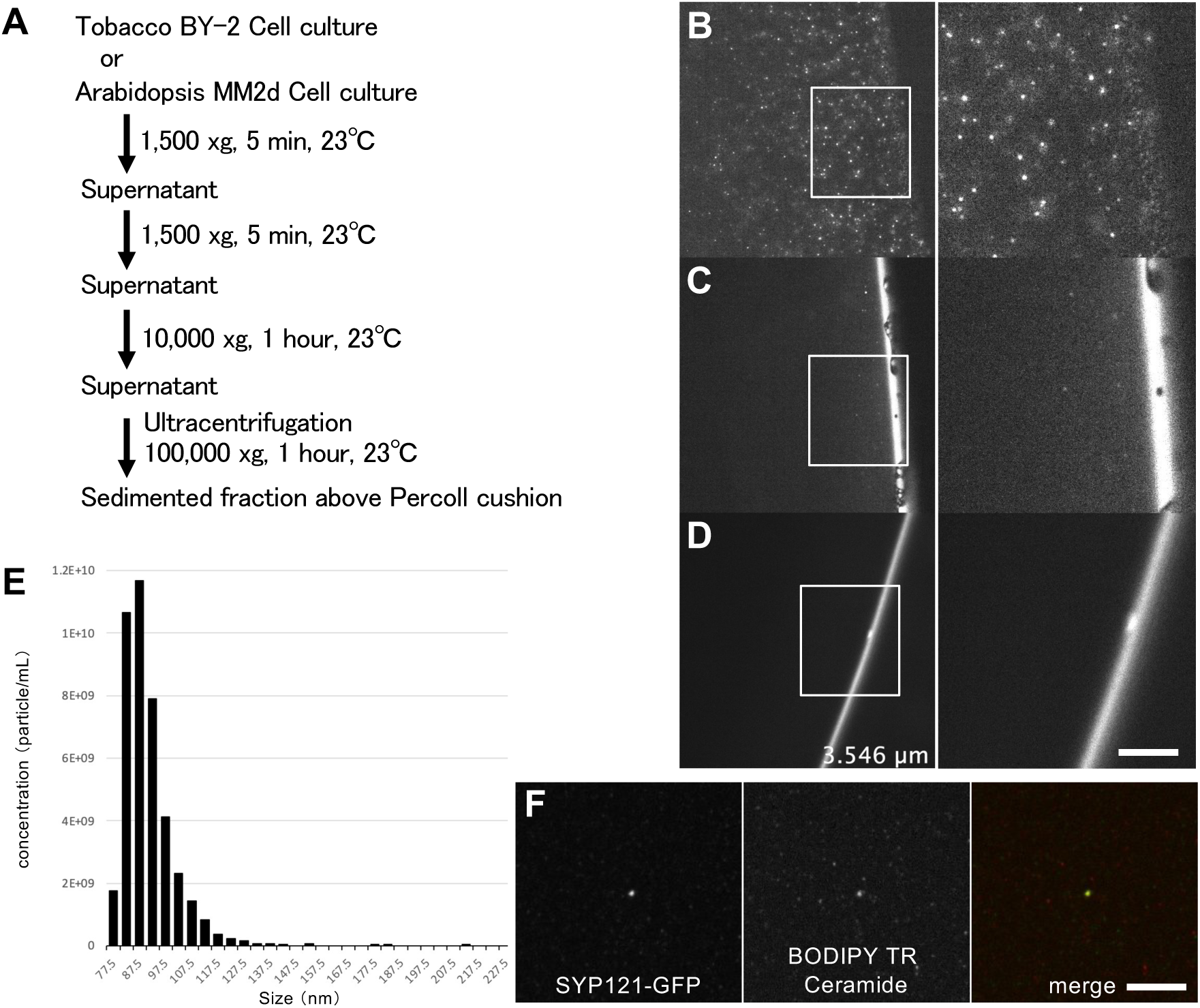
Isolation of EVs from a plant cell culture suspension. **A)** Protocol for the isolation of EVs from plant cell cultures. **B–D)** BODIPY TR Ceramide staining of the ultracentrifugation-derived sedimented fraction (B) and supernatant fraction (C) for the tobacco BY-2 culture medium and the ultracentrifugation-derived sedimented fraction for the medium before culturing (D). The panels on the right present the enlarged images of the area marked by a white square in the panels on the left. These images correspond to the optical section presented in Supplemental Movie 1. Bar, 2 µm. **E)** Size distribution of sedimented particles by ultracentrifugation from the supernatant of Arabidopsis MM2d cells. **F)** Purified EVs from SYP121-GFP transgenic tobacco BY-2 cell cultures. Bar, 2 µm.

The Arabidopsis cell culture EVs proteome revealed the EV protein composition more accurately than the earlier seedling apoplast EV proteomes

To characterize the cell culture EVs, we collected EVs from the supernatant of a healthy 7-day old MM2d cell culture (Supplemental Figure 1) and analyzed the protein composition of the Arabidopsis MM2d EV fraction by mass spectrometry. A total of 1,356 proteins were identified in the cell culture EV proteome (Supplemental Table 1), including 8.2% and 24.1% of the proteins in the seedling apoplast EV proteome identified by He et al. (2021) and Rutter and Innes (2017), respectively (Fig. 2A). The major proteins in the cell culture EV proteome were apoplast-related proteins, membrane trafficking proteins, cell wall-associated proteins, plasma membrane proteins, membrane transporters, and PD-related proteins (Fig. 2B).

**Figure 2.**
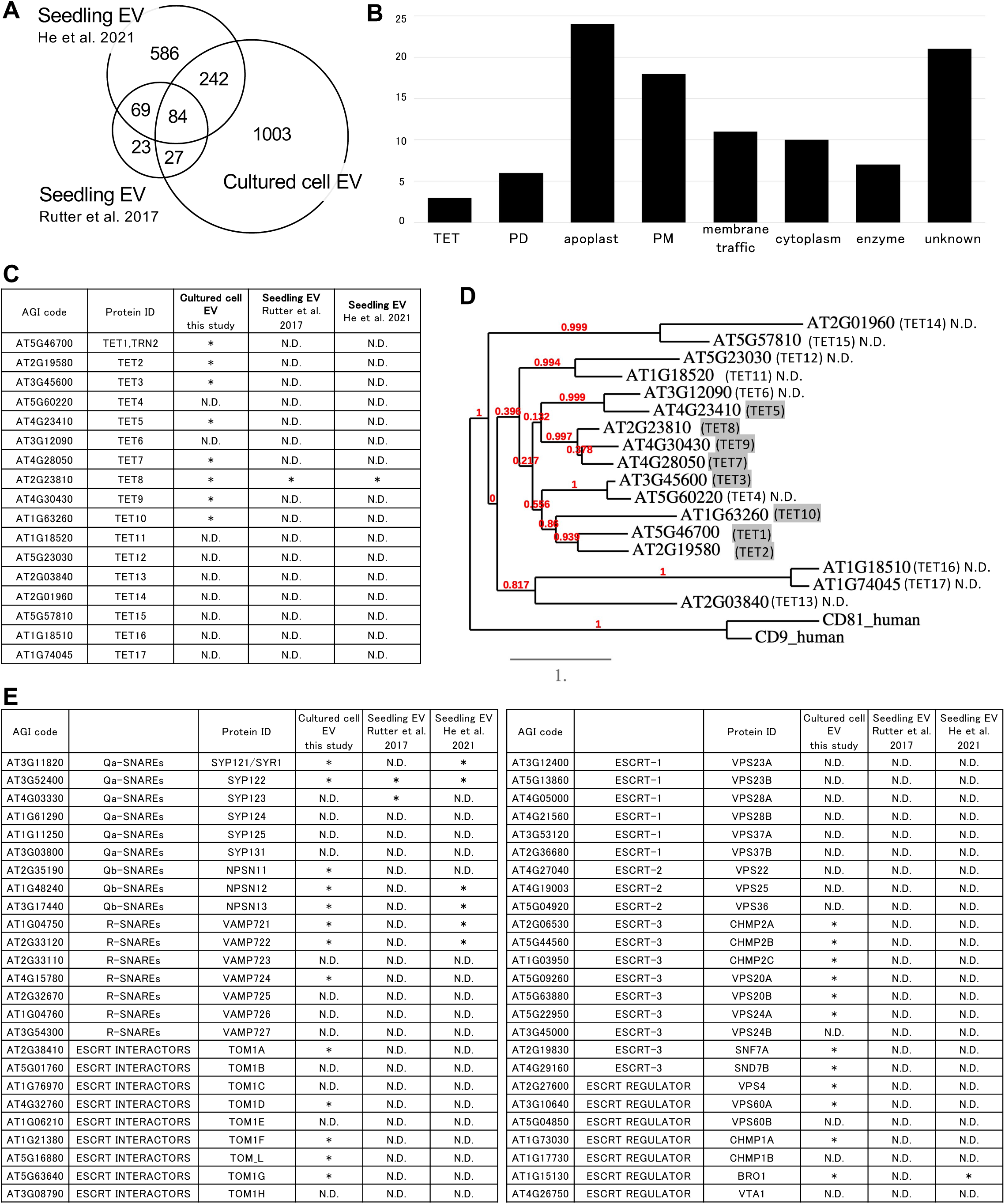
Comparison between the cell culture EV proteome and the seedling apoplast EV proteome. **A)** Venn diagram of the identified proteins in the cell culture EV proteome and the seedling apoplast EV proteome. **B)** Classification of the top 100 high-intensity proteins in the cell culture EV proteome. **C)** Identified tetraspanin family members in the cell culture EV proteome and the seedling apoplast EV proteome. **D)** Phylogenetic tree of the tetraspanin family in *Arabidopsis thaliana*. Identified tetraspanin family members in the cell culture EV proteome were colored in gray. **E)** Identified membrane trafficking proteins in the cell culture EV proteome and the seedling apoplast EV proteome.

Compared with the earlier analyses of seedling apoplast EV proteomes, our examination of the cell culture EV proteome identified more proteins, including specific protein subgroups predicted to have overlapping functions. This indicates that cell culture EVs may provide a more comprehensive overview of the EV proteome than seedling apoplast EVs. For example, of the 17 tetraspanin proteins in Arabidopsis (Cnops et al., 2006), only one (TET8) was identified in the seedling apoplast EV proteome (Rutter and Innes, 2017; He et al., 2021) (Fig. 2C). In contrast, eight of the 10 highly conserved plant tetraspanins (TET1–TET10) were identified in the cell culture EVs (Fig. 2D). TET11– TET17, which are less conserved tetraspanin family members (Fig. 2D) and lack homologs in other plants (Cnops et al., 2006), were not identified. Accordingly, TET1– TET10 are likely the functional tetraspanins in plant EVs.

Membrane trafficking proteins were also identified as EV proteins (Fig. 2E). For example, SYP121 and its functional ortholog SYP122, were identified in both the cell culture EV and seedling apoplast EV proteomes. In addition, VAMP721/VAMP722, which participate in the fusion of MVBs to the plasma membrane together with SYP121/SYP122 (Kwon et al., 2008), were also identified in both the cell culture EV and seedling apoplast EV proteomes. Unlike seedling apoplast EV proteomes, the cell culture EV proteome was enriched with specific subgroups of MVBs, such as ESCRT-0 and ESCRT-3 complexes and the VPS4 complex, but not ESCRT-1 and ESCRT-2 (Fig. 2E). Thus, compared with the previously characterized EV proteomes, the cell culture EV proteome may provide additional insights into plant EV protein compositions.

In addition to the typical EV proteins, the cell culture EV proteome contained many membrane trafficking proteins, including specific subgroups of small GTPases (ARF, RAB, and ROP) (Supplemental Table 2). Earlier research showed ARA6/AtRABF1 helps mediate the fusion of MVBs to plasma membranes (Ebine et al., 2012). Additionally, AtRABA1a is colocalized with VAMP721/722 and facilitates the transport between the TGN and the plasma membrane (Asaoka et al., 2013). Both AtRABD2a and AtRABD2b are also involved in the transport from the TGN to the plasma membrane (Pinheiro et al., 2009; Drakakaki et al., 2012). The SCAMP family members, which help transport vesicles to the plasma membrane (Toyooka et al., 2009), were also enriched in the cell culture EV proteome. There were some differences between the cell culture EV proteome and the seedling apoplast EV proteome. More specifically, all clathrin heavy and light chains and many clathrin-related proteins were detected in the seedling apoplast EV proteome (He et al., 2021). Conversely, most of these proteins were not identified in the cell culture EV proteome (Supplemental Table 2). Similarly, there were differences in the identified EXOCYST-related components (Supplemental Table 2). The membrane trafficking proteins detected in the cell culture EV proteome mediate various types of transport to the plasma membrane, suggesting the cell culture EV fraction contained various types of EVs.

Transporters, proton pumps, and auxin carriers, which are plasma membrane proteins, were also detected in the cell culture EV proteome (Supplemental Table 3). These proteins are ubiquitinated and transported to vacuoles for degradation via the MVB pathway (Mosesso et al., 2019). If MVBs are transported to the plasma membrane rather than to the vacuolar membrane, the vesicles within the MVBs are considered to be EVs. Therefore, it is not surprising that these proteins were identified in EVs.

O-Glycosyl hydrolases that degrade β-1,3-glucan were relatively abundant in the cell culture EV fraction. Among the 51 O-glycosyl hydrolase family 17 members in Arabidopsis, 25 were previously predicted to contain the GPI-anchor motif that binds to membranes (Doxey et al., 2007). In the cell culture EV proteome, 23 O-glycosyl hydrolases were identified, of which 18 had the GPI-anchor motif, suggesting that membrane-bound O-glycosyl hydrolases are selectively concentrated in the cell culture EV proteome (Supplemental Table 4). Some O-glycosyl hydrolases were detected in seedling apoplast EV proteomes, but they have not been thoroughly characterized.

The cell culture EV proteome also included several enzymes, such as strictosidine synthase, eukaryotic aspartyl protease, purple acid phosphatase, and rotamase CYP, which were previously identified in the apoplast proteome (Supplemental Table 1) (Uemura et al., 2019). Although their exact functions are unknown, these enzymes might modulate secondary metabolite metabolism or cell wall regulation in apoplasts.

### The EV and PD proteomes had similar compositions

Rutter and Innes (2017) reported that the identified proteins in the seedling apoplast EV proteome overlapped approximately 50% of the identified proteins in the PD proteome (Fernandez-Calvino et al., 2011) and the membrane trafficking-related proteome (Heard et al., 2015). In the current study, the identified proteins in the cell culture EV proteome included many PD proteins. In terms of the top 30 high-intensity proteins in the cell culture EV proteome, major plant EV markers, such as tetraspanins (TET8 and TET1) and SNAREs (SYP121, SYP122, and SYP132), were identified in both EV and PD proteomes (Fig. 3A) (Fernandez-Calvino et al., 2011; Brault et al., 2019). Moreover, major PD markers, such as PDLP and PCBP family members, were also identified in the cell culture EV proteome (Fig. 3A). Several specific proteins, including the disease resistance-related protein NDR1, late embryogenesis abundant (LEA) proteins, O-glycosyl hydrolases, and oxidases (including SKU5 family members) were common to the EV and PD proteomes (Fig. 3A).

**Figure 3.**
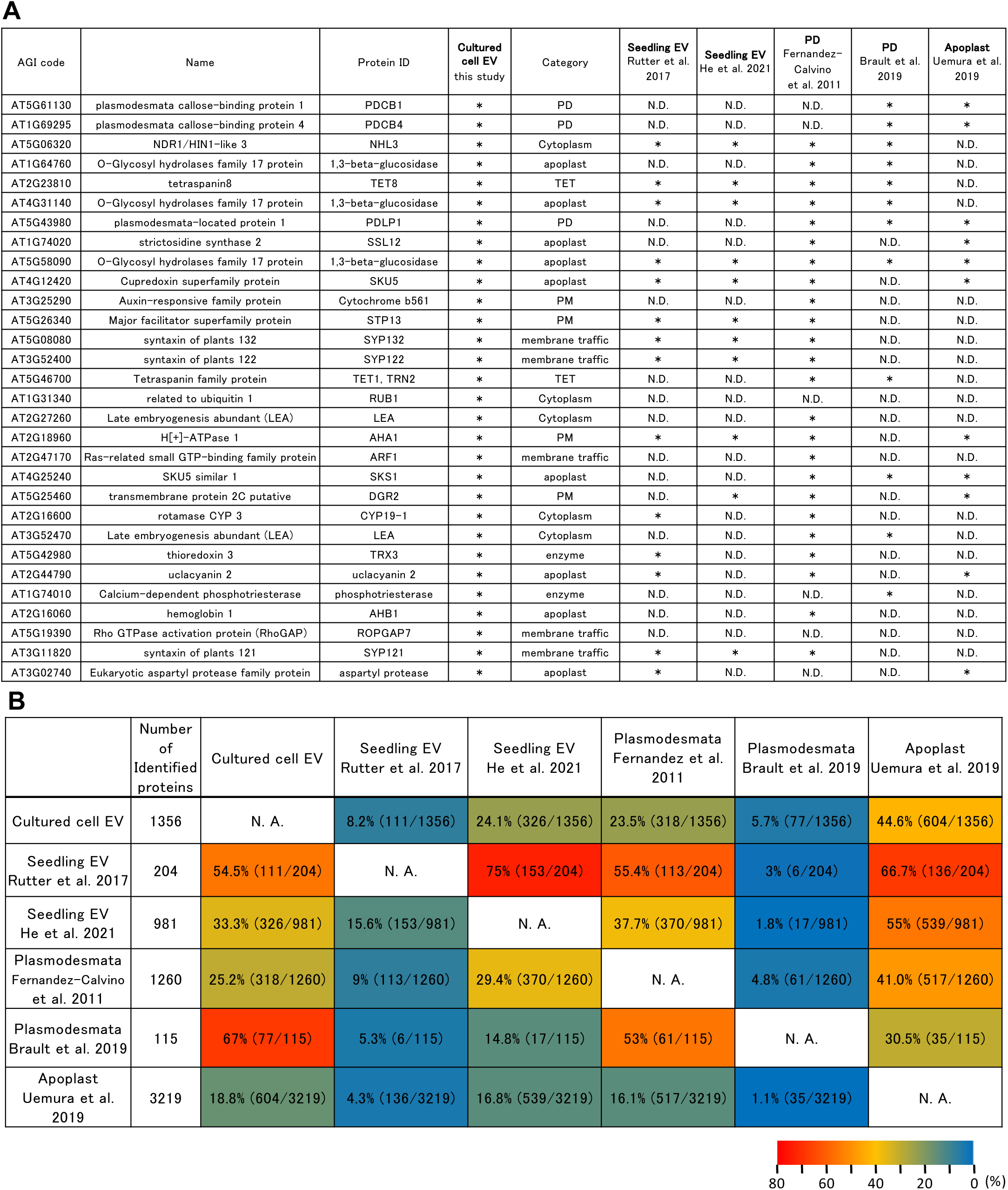
Comparison between the EV proteome and the plasmodesmata proteome. **A)** Comparison of the EV proteomes and the plasmodesmata proteomes in terms of the top 30 high-intensity proteins in the cell culture EV proteomes. **B)** Overlap percentages between the three EV proteomes and the two plasmodesmata proteomes. The EV and plasmodesmata fractions were indistinguishable in these analyses.

Of the 115 proteins identified in an earlier analysis of a PD proteome (Brault et al., 2019), 77 were detected in the cell culture EV proteome (Fig. 3B). Among the 50 proteins with the highest scores in the PD proteome (Brault et al., 2019), 43 were also identified in the cell culture EVs. A similar correlation was also observed between the seedling apoplast EV proteome (Rutter and Innes, 2017) and the PD proteome (Fernandez-Calvino et al., 2011); 113 proteins in the PD proteome were also among the 204 proteins in the seedling EV proteome (Fig. 3B). In the previous proteomic analyses, PDs were isolated from the supernatant of cell wall-digested cultures by differential centrifugation. Isolated PDs are reported as membranous components containing 50–100 nm vesicle-like structures (Fernandez-Calvino et al., 2011). Hence, they are similar in size to EVs. Due to the similarities in the purification protocols and vesicle sizes, it seems that the separation of EVs from PDs was difficult.

### PDLP3 localized to small particles in the apoplast and was distinguishable from EVs

This study identified the amount of PD-related factors in the cultured cell supernatant fraction (Fig 3). However, large membrane structures, as seen in the purified fraction in the PD proteome analysis (Fernandez-Calvino et al., 2011), were not observed in the cultured cell supernatant fraction. Therefore, we considered the possibility that PD-related factors are secreted into the extracellular space as small particles.

To assess this possibility, we used an established PD marker line, pPDLP3::PDLP3-GFP transgenic seedling (Thomas et al., 2008) and checked the localization by a spinning-disc confocal laser microscopy equipped with hyper-sensitive sCMOS cameras (Fig. 4A). PDLP3-GFP clearly localized to PDs (Fig. 4A). In addition, observations with a hyper-sensitive camera showed that PDLP3-GFP localized to particles in cells such as membrane trafficking vesicles (Fig. 4B) as previously reported (Thomas et al., 2008). Observations with a hyper-sensitive camera also revealed that PDLP3-GFP particles localized to both the plasma membrane and the extracellular region of hypocotyl epidermal cells (Fig. 4C, D). Timelapse analyses revealed that signals of extracellular PDLP3-GFP particles were immobile, sticking one spot for a long time (Fig. 4E, Supplemental Movie 2), whereas signals of plasma membrane-bound PDLP3-GFP particles showed intense two-dimensional motility (Fig. 4F, Supplemental Movie 2). These results showed that PDLP3-GFP secreted into apoplastic regions where PDs are absent.

**Figure 4.**
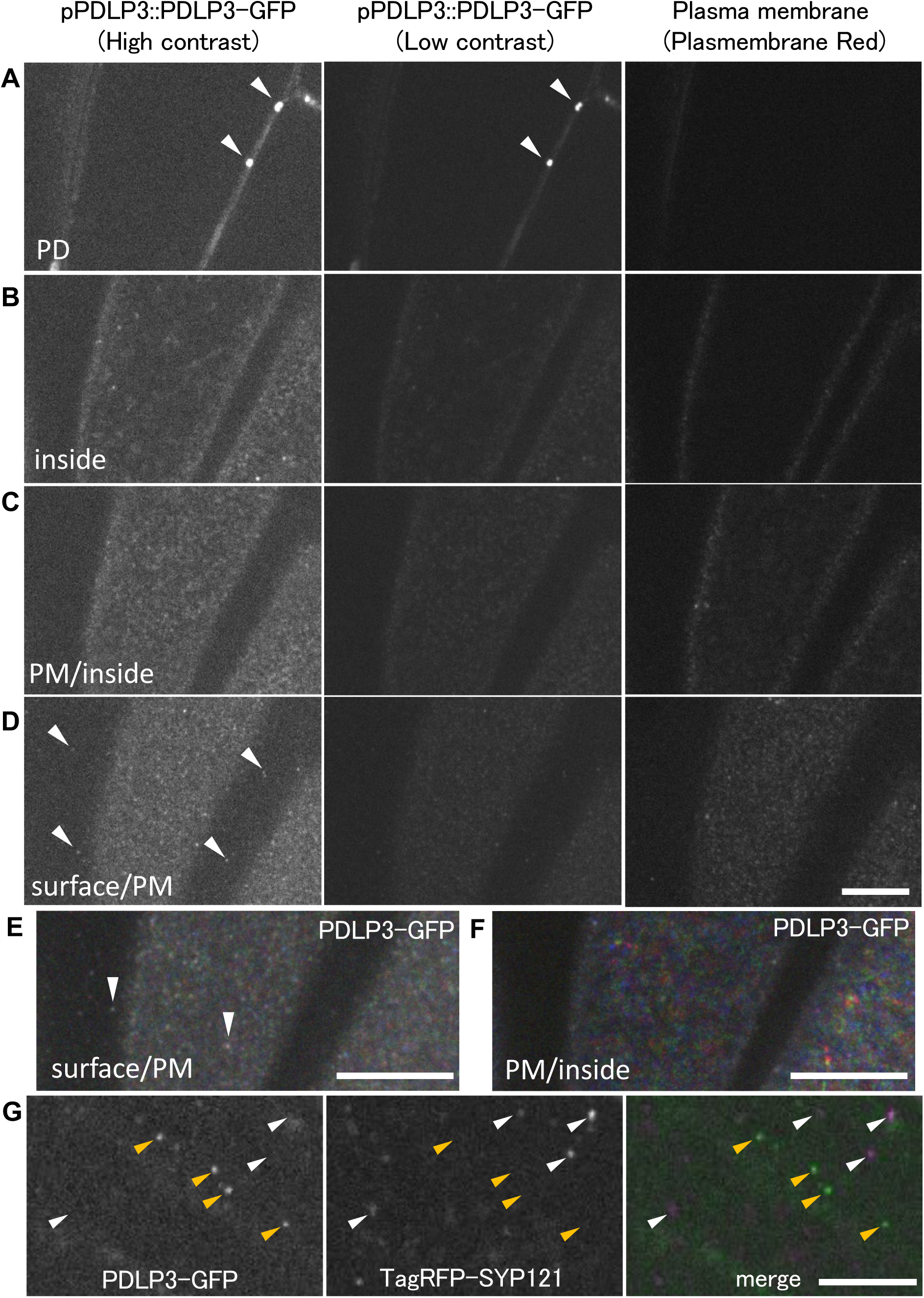
Localization of PDLP3-GFP in the apoplast region. **A-D)** Confocal microscopy images of PDLP3-GFP. Plasma membrane was labeled by Plasmembrane Red. PDLP3-GFP clearly localized to plasmodesmata (A, white arrowheads). At high contrast images, the PDLP3-GFP signals were also observed as particles at inside (B), plasma membrane (C), and surface (D). Some PDLP3-GFP signals were detected as small particles in the apoplast region (D, white arrowheads). Bar, 10 µm. **E, F)** Projection images from PDLP3-GFP time-lapse analyses. The 15 time-lapse images were acquired every 5 seconds and grouped into early (1-5), middle (6-10), and late (11-15) phases. These images were projected at maximum intensity, and superimposed into one merge image using blue, green and red colour, respectively. Bars, 10 µm. **G)** Comparison of PDLP3-GFP and TagRFP-SYP121 at the apoplast region. TagRFP-SYP121 (white arrowheads) were not colocalized with PDLP3-GFP (yellow arrowheads). Bar, 5 µm.

Next, we checked whether the extracellular PDLP3-GFP particles were identical to EVs. To assess this possibility, we selected TagRFP-SYP121 (Ebine et al. 2011) as an EV marker. Observation of the sample mounting water of stable transgenic plants expressing PDLP3-GFP and TagRFP-SYP121 showed that the particles of PDLP3-GFP and TagRFP-SYP121 were independent (Fig. 4G), indicating that the extracellular PDLP3-GFP particles are not identical to EVs.

### The EV and PD proteins were distinguished by a density gradient ultracentrifugation

The differences in the EV and PD protein localization implied these proteins may be separated experimentally. To generate more pure EV fractions, we separated the crude EV fractions by density gradient ultracentrifugation. The crude EV fractions were collected from the cell culture supernatants by ultracentrifugation and then used for the 24%–3% step-wise iodixanol density gradient ultracentrifugation, which resulted in six fractions (Fig. 5A) that were subjected to the size distribution analysis and the proteome analyses. Although the size distribution analysis did not show significant differences between fractions (Supplemental Figure 2), the proteome analyses identified 524 proteins (Fig. 5B, Supplemental Table 5).

**Figure 5.**
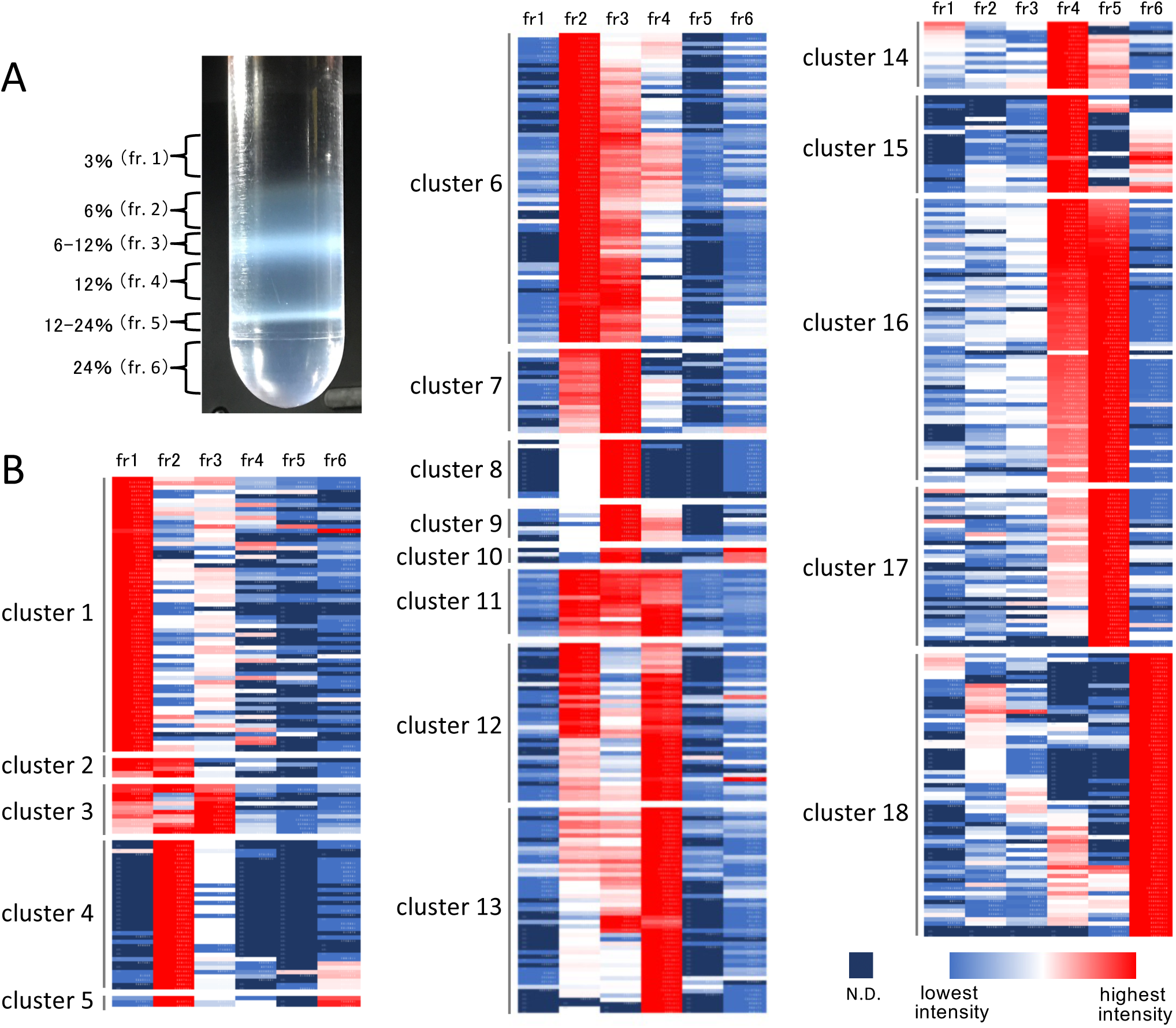
Iodixanol density gradient ultracentrifugation of the crude EV fraction. A) Separation of the crude EV fraction by an iodixanol density gradient ultracentrifugation. The crude EV fraction was mounted on the top of the iodixanol solution during the density gradient ultracentrifugation (24%, 12%, 6%, and 3% iodixanol solutions). After the ultracentrifugation, six fractions were recovered and used for proteome analyses. B) Classification of the 524 identified proteins on the basis of their distribution. The presented data correspond to the information provided in Supplemental Table 5.

The 524 identified proteins were classified into 18 clusters according to their distribution (Fig. 5B, Supplemental Table 5). Tetraspanin and SNARE proteins were distributed between 6% and 12% (fractions 2, 3, and 4), suggesting that typical EVs in plant cell cultures are distributed between 6% and 12% (Fig. 6A). The PDLP (PDLP1, 3, and 6) and PDCB (PDCB1 and 4) proteins, which are representative PD proteins, were also distributed between 6% and 12% (fractions 2, 3, and 4) (Fig. 6B). The EV markers were often most concentrated in fraction 4, followed by fractions 2 and then 3 or they were exclusively concentrated in fraction 4 (Fig. 6A), whereas the PD markers were most enriched in fraction 2, followed by fractions 3 and then 4 (Fig. 6B). Accordingly, although it is likely difficult to completely separate EVs and PD, the EV markers can be distinguished from the PD markers by iodixanol density gradient ultracentrifugation and clustering analyses.

**Figure 6.**
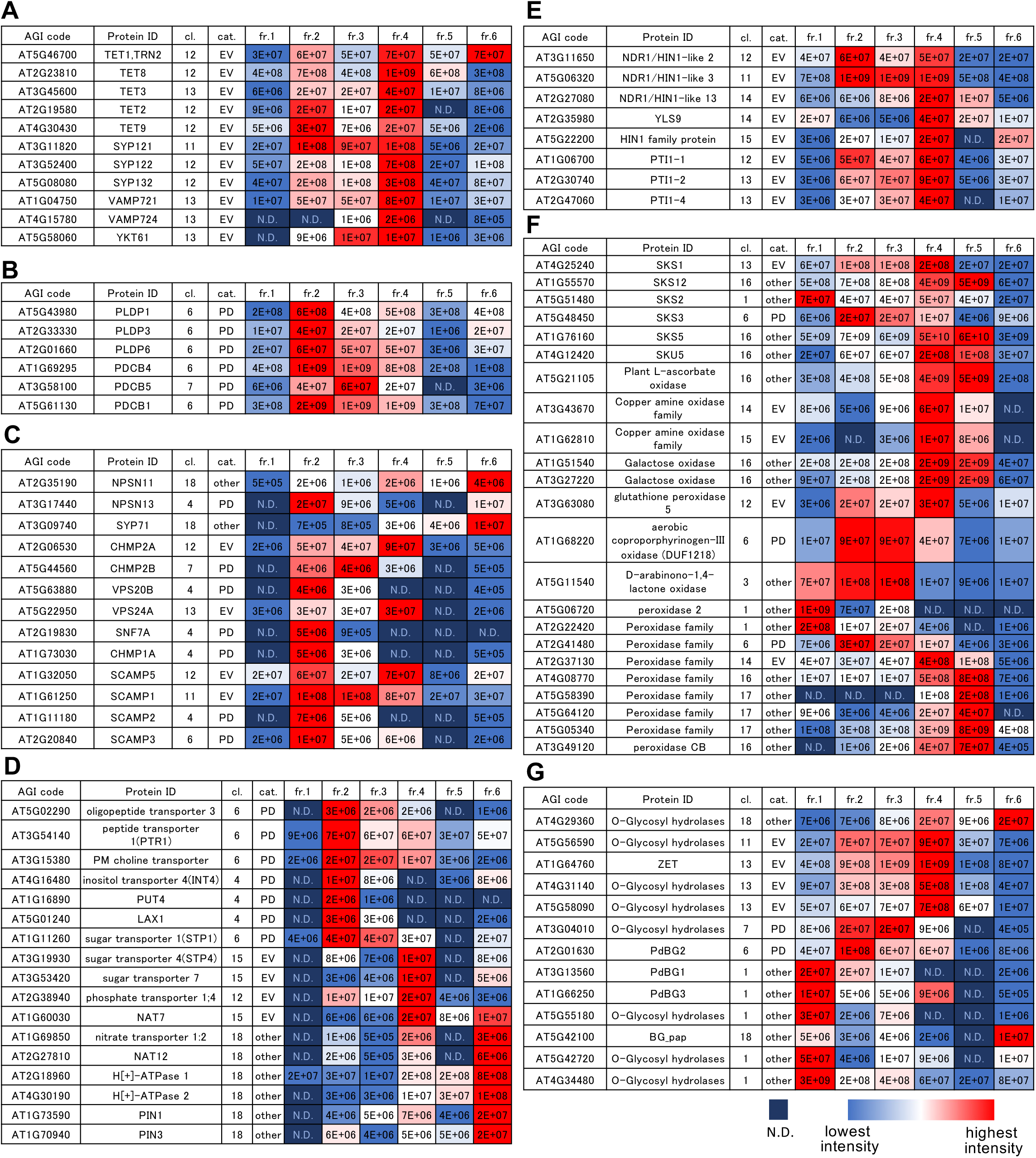
Representative proteins in each gene ontology category. A) Representative EV proteins. B) Representative PD proteins. C) Membrane trafficking-related proteins. D) Membrane transporters and pumps. E) Disease resistance-related proteins. F) Oxidases and peroxidases. G) O-glycosyl hydrolases. The presented data correspond to the information provided in Supplemental Table 5. cl. indicates the cluster number in Figure 5B; cat. indicates category.

We subsequently classified additional proteins in the EV, PD, and Other categories. Among the membrane trafficking markers, the SNAREs VAMP721, VAMP724, and YKT61 as well as SYP121 and SYP122 were classified in the EV category (Fig. 6A). Among the ESCRT proteins involved in MVB formation, CHMP2A and VPS24A were classified in the EV category, whereas CHMP2B, VPS20B, and SNF7A were classified in the PD category (Fig. 6C). Moreover, SCAMP and the RAB GTPases were also divided into the EV and PD categories (Fig. 6C). The crude EV fraction contained many transporters belonging to the PD, EV, and Other categories (Fig. 6D). In the Other category, NRT1:2, proton transporters, and auxin transporters were concentrated in the heaviest fraction (i.e., fraction 6). The NDR1/HIN1 family, which is involved in disease resistance, was clearly concentrated in the EV category (Fig. 6E). The LEA family, which is phylogenetically closely related to the NDR1/HIN1 family, was classified in either the EV or PD category (Supplemental Table 5). In addition, the PTI1 family, which comprises key signaling proteins related to disease resistance (Sun et al., 2022), was concentrated in the EV category (Fig. 6E).

A number of transporters were classified in the PD category, among which CHER1 (choline transporter) is localized in TGNs, cell plates, and sieve pores after cell division; it is important for the development of sieve pores and PD (Dettmer et al., 2014; Kraner et al., 2017; Gao et al., 2017b). Although CHER1 has not been confirmed as a PD-localized protein, it was classified in the PD category in this study. Other transporters in the PD category may also affect PD development and functions.

Oxidases and peroxidases, which regulate the apoplast redox state and pH homeostasis, were relatively abundant in the crude EV fraction. The apoplast redox state and pH homeostasis are crucial factors influencing cell expansion, development, and defenses against pathogens (Smirnoff and Arnaud, 2019). Both SKU5 and SKS1 contribute to cell wall formation (Chen et al., 2023; Sedbrook et al., 2002). In the present study, SKS3 and SKS1 were classified in the PD and EV categories, respectively (Fig. 6F). Furthermore, SKU5, SKS5, and SKS12 were concentrated in fractions 4 and 5, which were slightly heavier than the EV fraction. Interestingly, most of the other oxidases and peroxidases were also concentrated in fractions 4 and 5. Beta-glucosidases were also enriched in fractions 4 and 5, one of which (BGLU15) is an apoplastic enzyme that mediates the degradation of flavonol bisglycosides (Roepke et al., 2017). These results suggest that oxidases, peroxidases, and beta-glucosidases are localized to some apoplastic structures other than EVs and PD.

The beta-1,3-glucanases, which are O-glycosyl hydrolases involved in degrading callose, were widely distributed in each fraction. Earlier research showed PdBG1, PdBG2, and PdBG3 are localized to PD (Levy et al., 2007; Benitez-Alfonso et al., 2013). In the current study, PdBG2 was classified in the PD category, but PdBG1 and PdBG3 were concentrated in the lightest fraction (i.e., fraction 1) (Fig. 6G). Moreover, BG_pap is a PD-localized beta-1,3-glucanase (Levy et al., 2007; Zavaliev et al., 2016). However, BG_pap was abundant in the heaviest fraction (i.e., fraction 6) (Fig. 6G). The EV fractions contained ZERZAUST, which is an important O-glycosyl hydrolase for intercellular communication (Vaddepalli et al., 2017). In addition, several O-glycosyl hydrolases were enriched in the EV fraction (Fig. 6G). These findings suggest that O-glycosyl hydrolases are localized to various structures in the apoplast.

## Discussion

Extracellular vesicles are membrane-coated vesicles in apoplasts. These vesicles are transported via various membrane trafficking pathways and degradation pathways. Plasmodesmata connect neighboring cells as cell wall-embedded structures consisting of the plasma membrane, cytoplasm, and ER. The apoplast side of PD accumulates callose and some specific components, including callose-binding proteins, callose synthases, and callose degrading enzymes (β-1,3-glucanases). On the basis of these distinct structural differences, EVs and PD are generally considered to be different structures.

The EV proteomes analyzed in a previous study (Rutter and Innes, 2017) and in the current study were revealed to contain a considerable abundance of PD proteins. Similarly, earlier examinations of PD proteomes identified EV proteins (Fernandez-Calvino et al., 2011; Brault et al., 2019). Hence, EV and PD proteins were not separated in previous proteome-level investigations, possibly because of the similarities in the purification methods. More specifically, EVs were purified from the apoplast fraction by centrifugation, while PD were purified via the centrifugation of solutions comprising digested cell walls. The reported size of EVs (50–300 nm) (Rutter and Innes, 2017) is similar to the size of vesicle-like PD structures purified from the cellulase-digested cell wall fraction (50–100 nm) (Fernandez-Calvino et al., 2011).

In the present study, we distinguished EVs from PD on the basis of a density gradient centrifugation and fine-tuned clustering. We revealed the differences in the clustering of typical EV markers, such as tetraspanins and specific SNARE family members, and typical PD markers, such as PDLP and PDCB proteins. Recent analyses of the *Physcomitrium patens* PD proteome successfully identified core PD proteins (Johnston et al., 2023; Gombos et al., 2023). The PDLP and PDCB families are apparently not conserved in *P. patens* or in *Marchantia polymorpha* and *Klebsormidium nitens* (Bowman et al., 2017), implying that the fraction containing the PDLP and PDCB proteins in this study may not be the PD fraction that includes the desmotubule. A number of unclassified clusters were identified in this study (Figure 5 and Supplemental Table 5). Of these, cluster 18, which was enriched in the heaviest fraction (i.e., fraction 6), is notable because it contained MCTP4, but lacked typical PD markers. The MCTP4 protein belongs to the MCTP family, which may be involved in linking the ER membrane and the plasma membrane within PD (Brault et al., 2019). Additionally, among the SNARE proteins, only SYP71 was classified in cluster 18. This protein is localized to the plasma membrane, ER, and cell plate (Suwastika et al., 2008; El Kasmi et al., 2013). These results suggest that cluster 18 may be enriched with proteins linking the desmotubule to the plasma membrane in PD. In the current study, the PD fraction including the PDLP and PDCB proteins was slightly lighter than the EV fraction, suggesting that the PD fraction may consist of only small particles (e.g., vesicles) and proteins that aggregate on the apoplast side of PD. The heavy fraction (i.e., cluster 18) may contain PD. Clarifying whether the proteins in cluster 18 actually belong to desmotubule–plasma membrane structures in PD may require additional electron microscopy analyses.

In contrast to plant seedlings, cultured cells are undifferentiated and homogeneous. In this study, we originally planned to recover pure homogeneous EVs via the ultracentrifugation of a large volume of cell culture supernatant. However, the recovered fraction contained various apoplast granules, including PD-associated proteins and apoplast proteins as well as EV proteins. The density gradient centrifugation and clustering categorized these proteins into 18 clusters, but it is unclear whether apoplast granules and structures remained. The study findings indicate that the secretion of apoplast materials is a complex process, even in homogeneous cell cultures. Plant seedlings consisting of various differentiated cells secrete more complex apoplast components. Furthermore, the types and amount of secreted apoplast components are influenced by biotic and abiotic stimuli. Future analyses of the clusters identified in this study will further characterize apoplast structures and physiological responses in plant seedlings.

## Materials and Methods

### Growth conditions of Arabidopsis cultured cells

Arabidopsis MM2d cells (Menges and Murray, 2002) and tobacco BY-2 cells were cultured in modified LS medium (Nagata et al., 1981) on a rotary shaker (130 rpm) at 27 °C in darkness. To maintain the cell culture, 3 mL 7-day-old MM2d cultured cells were transferred to 95 mL fresh medium in 300 mL flasks. Arabidopsis plants (Col-0) were grown on 1.5% agar medium containing 1% sucrose and 1% MES-KOH (pH 5.7) in plates and in half-strength Murashige and Skoog salt medium at 22 °C for 5 days.

### Isolation of EVs from the culture medium

Seven-day-old MM2d cell cultures were centrifuged twice (1,500 × g for 5 min). The supernatants were collected for the high-speed centrifugation (10,000 × g for 1 h), which was completed to eliminate any remaining debris. The purified cell culture supernatants were placed on a Percoll cushion (37% Percoll and 0.6 M sorbitol) prepared in a centrifuge tube for the subsequent ultracentrifugation (100,000 × g for 1 h). The white layer on the Percoll cushion was recovered. The EVs were further purified using exoEasy Maxi kit (Qiagen). The resulting samples, which were designated as EV fractions, were used for the mass spectrometry analysis.

For the density gradient ultracentrifugation, the supernatants of 5-day-old MM2d cell cultures were prepared via the centrifugations described above and collected from the top of the 24% iodixanol solution containing 0.25 M sucrose and 20 mM HEPES-KOH (pH 7.4) after the ultracentrifugation (100,000 × g for 1 h). The collected crude EV fractions were diluted with 10 volumes of buffer (0.25 M sucrose and 20 mM HEPES-KOH, pH 7.4) before the density gradient ultracentrifugation (100,000 × g for 1 h) involving the 24%, 12%, 6%, and 3% step-wise iodixanol density gradient. Each layer was recovered and used for the mass spectrometry analysis.

### Size distribution and particle concentration

The size and number of particles in each fraction were analyzed using qNano (Izon, Christchurch, New Zealand). The NP200 nanopore and CPC100 calibration sample were used for particle detection and evaluation.

### Mass spectrometry

The mass spectrometry analysis was performed as described by Imami et al. (2018), with slight modifications. Proteins were precipitated in 20% trichloroacetic acid (TFA). The pelleted proteins were washed three times in ice-cold 100% acetone and then resuspended in 50 μL resuspension buffer (8 M urea and 0.1 M Tris-HCl, pH 8). After adding 10 mM dithiothreitol, the proteins were reduced at room temperature for 30 min and then alkylated using 50 mM iodoacetamide at room temperature for 30 min in darkness. Proteins were digested with lysyl endopeptidase (LysC) (Wako) at room temperature for 3 h. The sample solutions were diluted to a final concentration of 2 M urea and 50 mM ammonium bicarbonate prior to the trypsin (Promega) digestion, which was performed with constant agitation at room temperature for 16 h. The resulting peptide samples were desalted using SDB-XC (upper)/SCX (bottom) (GL Sciences) StageTips (Adachi et al., 2016). The peptides were sequentially eluted using elution buffer 1 (4% TFA, 500 mM ammonium acetate, and 30% acetonitrile) and elution buffer 2 (500 mM ammonium acetate and 30% acetonitrile).

The nano-scale reversed-phase liquid chromatography and tandem mass spectrometry (nanoLC/MS/MS) analysis was completed using the Orbitrap Fusion Lumos mass spectrometer (Thermo Fisher Scientific) connected to a Thermo Ultimate 3000 RSLCnano pump and an HTC-PAL autosampler (CTC Analytics, Zwingen, Switzerland) equipped with a self-packing analytical column (150 mm long with 100 μm internal diameter) (Ishihama et al., 2002) packed with ReproSil-Pur C18-AQ (3 μm; Dr. Maisch GmbH). Mobile phase A consisted of 0.5% acetic acid, whereas mobile phase B comprised 0.5% acetic acid and 80% acetonitrile. Peptides were eluted from the analytical column at a flow rate of 500 nL/min with the following gradient: 5%–10% B in 5 min, 10%–40% B in 20 min, 40%–99% B in 1 min, and 99% for 5 min. The Orbitrap Fusion Lumos instrument was operated in the data-dependent mode with a full scan in the Orbitrap followed by MS/MS scans for 3 s for the higher-energy collisional dissociation (HCD). The applied voltage for ionization was 2.4 kV. The full scans were performed with a resolution of 120,000, a target value of 4 × 10^5^ ions, and a maximum injection time of 50 ms. The MS scan range was m/z 300–1,500. The MS/MS scans were performed with a resolution of 15,000, a target value of 5 × 10^4^ ions, and a maximum injection time of 50 ms. The isolation window was set to 1.6. The normalized HCD collision energy was 30. Dynamic exclusion was applied for 20 s.

All raw data files were analyzed and processed using MaxQuant (v1.6.15.0) (Cox and Mann, 2008). Andromeda (Cox et al., 2011) was used to search the Araport11_genes.201606.pep (https://www.arabidopsis.org/) database spiked with common contaminants and enzyme sequences. The search parameters included two missed cleavage sites and variable modifications, including methionine oxidation, deamidation of glutamine and asparagine and protein N-terminal acetylation. Cysteine carbamidomethylation was set as a fixed modification. The peptide mass tolerance was 4.5 ppm, whereas the MS/MS tolerance was 20 ppm. The false discovery rate at the peptide spectrum match and protein levels was set to 1%.

### Microscopy analyses

The confocal microscopy analysis was performed using the ECLIPSE Ti microscope (Nikon) equipped with a confocal spinning disc unit (CSU-W1, Yokogawa) and the sCMOS camera (Sona, Andor). The isolated EVs were stained with 1 µM BODIPY TR Ceramide (ThermoFisher Scientific). For the optical sectioning of Arabidopsis seedlings, transgenic plants expressing both PDLP3-GFP and TagRFP-SYP121 were mounted between 60 × 24 and 18 × 18 coverslips (Iwanami) with a 0.15 mm spacer. Images were captured using NIS-Element AR (Nikon) and analyzed using Fiji (Schindelin et al., 2012).

## Acknowledgements and fundings

We thank Tsuyoshi Nakagawa (Shimane University) for providing Gateway binary vectors containing bar gene identified by Meiji Seika Kaisha, Ltd., Andrew J. Maule (John Innes Centre) for providing the PDLP3-GFP transgenic plant, Masa H. Sato and Tomoko Hirano (Kyoto Pref. Univ.) for providing the pGWB1-GFP-SYP121 plasmid, Kazuo Ebine and Takashi Ueda (NIBB) for providing the TagRFP-SYP121 seedling, Akihiko Nakano (Riken) and Tomohiro Uemura (Ochanimizu Univ) for helpful discussion. This study was supported by JST-PRESTO [JPMJPR18H7 to T. H., JPMJPR19H7 to K. Sueyoshi, and JPMJPR18H2 to K. I.] and JST-CREST [JPMJCR18H4 to T. H. and JPMJCR19H1 to K. Sueyoshi].

---

**Supplemental Figure 1.**
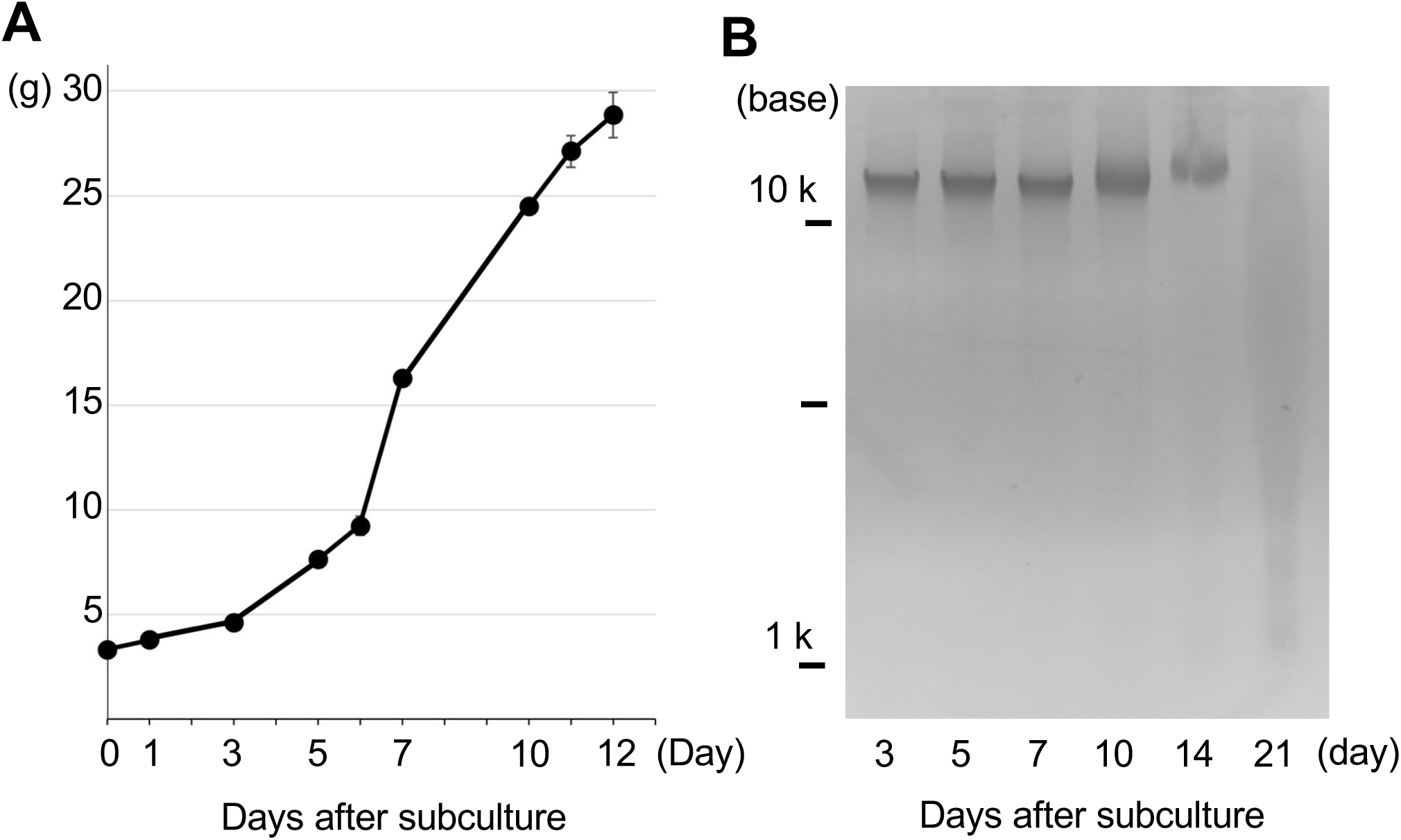
Growth condition of MM2d Arabidopsis cultured cells for this study. **A)** Growth curve of MM2d Arabidopsis cultured cells in this study. Supernatant of 5 days-old and 7 days-old cultured cells was applied for EV purification. **B)** DNA degradation assay of MM2d cells. In 5 days-old and 7 days-old cultured cells, DNA degradation did not occur at all.

**Supplemental Figure 2.**
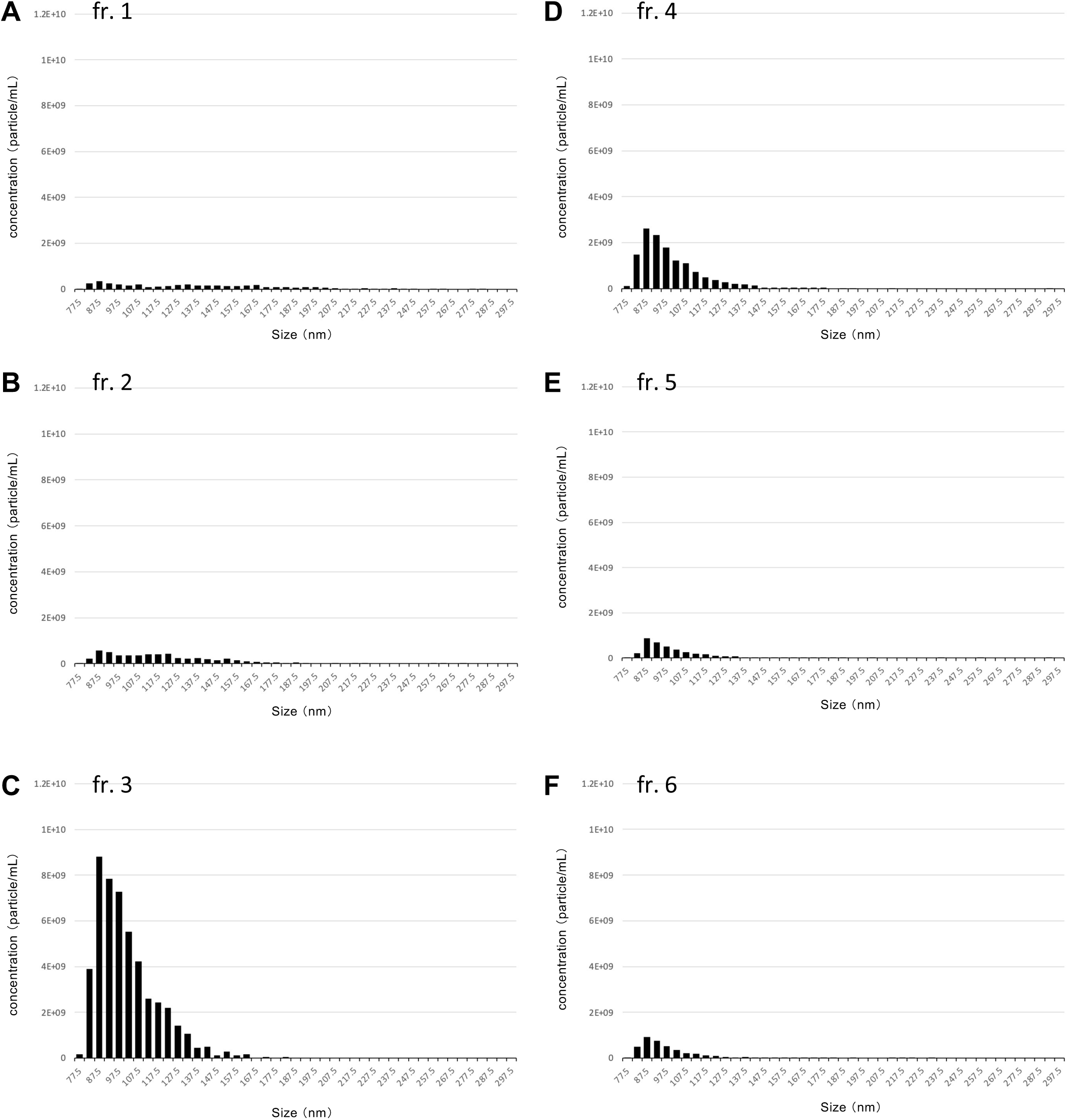
Size distribution of EVs in each fraction after an iodixanol density gradient ultracentrifugation. Six fractions were recovered after an iodixanol density gradient ultracentrifugation and used for size distribution analyses. These fractions correspond to the fractions in Figure 5A.

**Supplemental Table 1.** List of 1356 proteins identified in the cultured cell EV proteomics.

**Supplemental Table 2.** Comparison of membrane traffic proteins between the cultured cell and the leaf apolast EV proteomics.

**Supplemental Table 3.** Comparison of transpoters and ubiquitin-related proteins between the cultured cell and the leaf apolast EV proteomics.

**Supplemental Table 4.** Comparison of O-Glycosyl hydrolases family 17 proteins between the cultured cell and the leaf apolast EV proteomics.

**Supplemental Table 5.** List of 524 proteins Identified in the density gradient ultra-centrifugation proteome.

**Supplemental Movie 1 Optical sections of isolated EVs.**

EVs were stained by BODIPY TR Ceramide. Images were taken from the surface of coverslip toward to the top of a drop.

**Supplemental Movie 2 Time-lapse analyses of PDLP3-GFP.**

Images were taken from the same seedling at different depths. These images were used to generate the projection images of Figure 4E and F. Bars, 10 µm.

## Notes

### Competing Interest Statement

The authors have declared no competing interest.

### Summary of Updates

Correction of author name, partial correction of Abstract and Introduction

